# Alterations of the axon initial segment in multiple sclerosis

**DOI:** 10.1101/2022.03.07.483302

**Authors:** Aysegul Dilsizoglu Senol, Giulia Pinto, Maxime Beau, Vincent Guillemot, Jeff L. Dupree, Christine Stadelmann, Jonas Ranft, Catherine Lubetzki, Marc Davenne

## Abstract

Grey matter damage has been established as a key contributor to disability progression in multiple sclerosis. Aside from neuronal loss and axonal transections, which predominate in cortical demyelinated lesions, synaptic alterations have been detected in both demyelinated plaques and normal-appearing grey matter, resulting in functional neuronal damage. The axon initial segment is a key element of neuronal function, responsible for action potential initiation and maintenance of neuronal polarity. Despite several reports of profound axon initial segment alterations in different pathological models, among which experimental auto-immune encephalomyelitis, whether the axon initial segment is affected in multiple sclerosis is still unknown. Using immunohistochemistry, we analyzed axon initial segments from control and multiple sclerosis tissue, focusing on layer 5/6 pyramidal neurons in the neocortex and Purkinje cells in the cerebellum and performed analysis on the parameters known to control neuronal excitability, i.e., axon initial segment length and position. We found that the axon initial segment length was unchanged among the different multiple sclerosis samples, and not different from controls. In contrast, in both cell types, the axon initial segment position was altered, with an increased soma-axon initial segment gap, in both active and inactive demyelinated lesions. In addition, using a computational model, we show that this increased gap between soma and axon initial segment might increase neuronal excitability. Taken together, these results show for the first time changes of axon initial segments in multiple sclerosis, in active as well as inactive grey matter lesions in both neocortex and cerebellum, which might alter neuronal function.

## Introduction

It is now well established that in multiple sclerosis (MS), central nervous system pathology exceeds white matter, and that grey matter damage occurs early in disease evolution, correlating with clinical disability and cognitive dysfunction^1–4^. Associated with an inflammatory component consisting of both innate and adaptive immunity, grey matter damage combines structural and functional changes such as demyelination, neuritic transections and neuronal loss ^5,6^ as well as synaptic pathology ^7,8^. Whether the axon initial segment (AIS), a key player in neuronal function, is altered in MS grey matter is unknown.

Located next to the soma and immediately followed by the myelin sheath in most neurons, the AIS is the site of action potential (AP) initiation. This is due to the dense aggregation of voltage-gated sodium (Nav) channels by the cytoskeleton-linked anchoring protein, AnkyrinG (AnkG), which also clusters particular voltage-gated potassium (Kv) channels ^9^. This molecular architecture is very similar to that of nodes of Ranvier, which have been shown to be altered in MS white matter plaques ^10,11^. The AIS also acts as a barrier between the somatodendritic and axonal compartments due to an AIS-specific AnkG-organized cytoskeleton and plasma membrane composition, and as such maintains axonal integrity and neuronal polarity ^12,13^. AnkG is considered as the AIS master organizer protein and its expression along the AIS is thus critical for allowing the AIS to play its two key roles, spike initiation and maintenance of axonal identity. Indeed, AnkG loss leads to a total dismantling of AIS-constituent proteins, including Nav channels, failure of spike initiation and loss of axonal identity ^14–17^.

The AIS is a plastic domain: its length, its position relative to the soma and/or its ion channel composition can vary, depending for instance on the level of input activity received by the neuron ^18^. These AIS changes allow the neuron’s excitability properties to be modulated and in some cases homeostatically fine-tuned to the neuron’s environment ^18^). The AIS is also a vulnerable domain whose properties have been shown to be altered in many pathological conditions, as in Alzheimer’s disease models ^19–21^ in epileptic syndromes ^22^, ^23^, in an Angelman syndrome mouse model ^24^, or upon ischemia ^25^ or traumatic brain injury ^26^ in mice. To our knowledge AISs have not been analyzed in MS tissue, although AIS alterations have been reported in experimental models of MS. In the inflammatory myelin oligodendrocyte glycoprotein (MOG)-induced experimental allergic encephalomyelitis (EAE) mouse model, major AIS changes such as reduced length and AIS loss were found ^27^. In contrast, such changes were not detected in the (non-inflammatory) cuprizone-induced demyelination mouse model ^27,28^ where a change in AIS position (AIS onset closer to the soma) impacting excitability properties was reported. These experimental results paved the way for analyzing AISs in MS tissue. Here we show that AISs start further away from the soma in both cortical and cerebellar demyelinated grey matter, whereas their length remains unchanged. Furthermore, using a computational model, we provide evidence that this enlarged soma-AIS gap could potentially increase neuronal excitability.

## Materials and Methods

### Human post-mortem samples

Tissue samples and associated clinical and neuropathological data (Table 1) were supplied by the Multiple Sclerosis Society Tissue Bank funded by the Multiple Sclerosis Society of Great Britain and Northern Ireland, registered charity 207495. Despite many attempts, we were unable to visualize AISs (with antibodies directed against AIS proteins, such as AnkG or Nav) on paraffin-embedded sections. We therefore undertook our study on snap-frozen tissue. After post-mortem delays ranging from 5 to 24 hours, brains had been dissected out, immediately thereafter cut in 1cm-thick coronal slices then into 2×2×1cm^3^ blocks, before being frozen by immersion into −50°C isopentane and stored at −85°C.

**Table 1:**
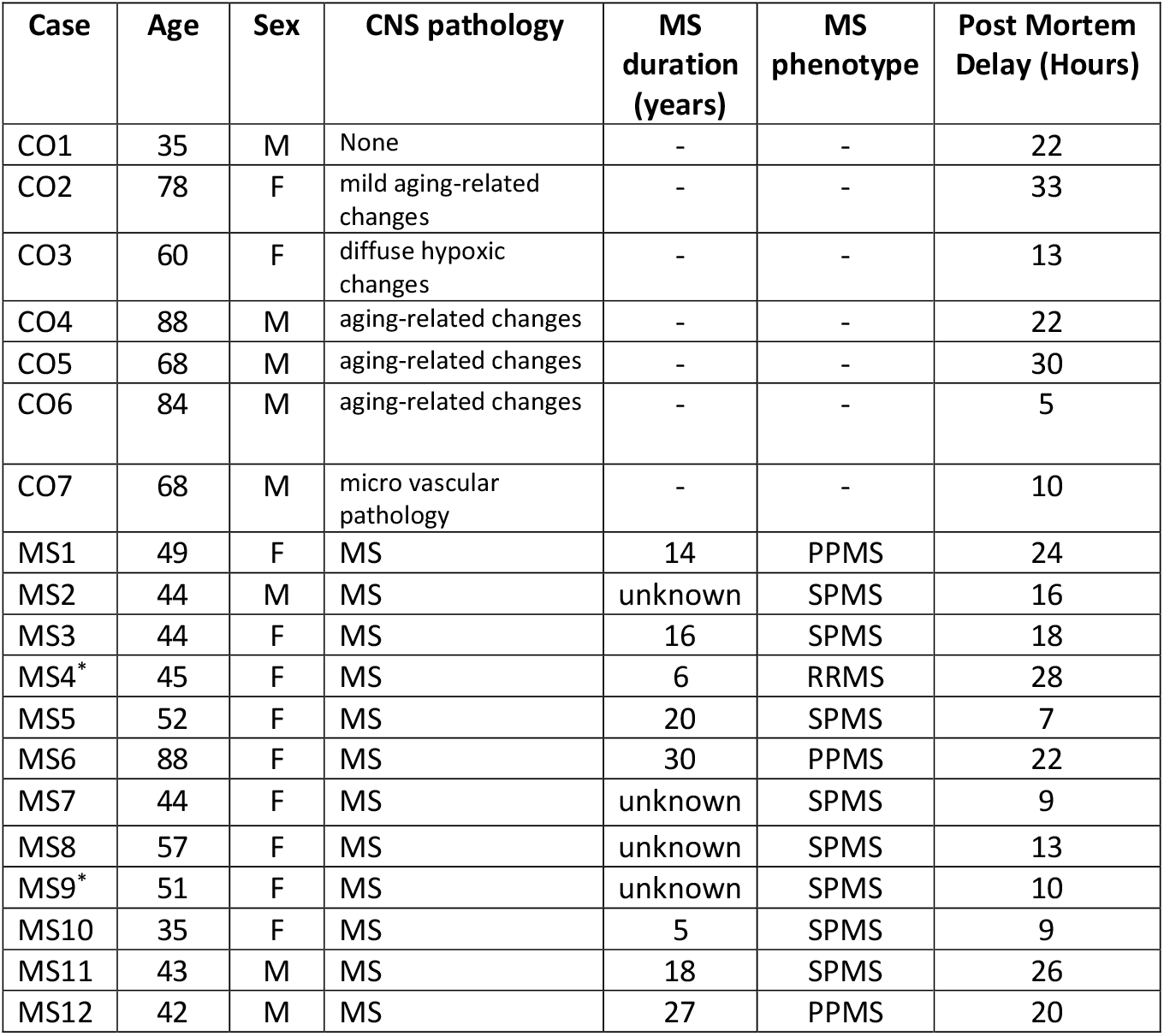
Patients characteristics. CO=control case; MS=Multiple sclerosis case; RRMS=relapsing-remitting MS; SPMS=secondary progressive MS; PPMS=primary progressive MS. * Normal appearing grey matter (NAGM), without cortical plaque (cerebellar sample only).

Table 1 shows the characteristics (age, sex, disease phenotype, disease duration) of the 12 MS cases and 7 control cases used in this study. As shown in Table 2, one to seven tissue blocks were available for each MS case. They consist of both neocortical and/or cerebellar cortical samples, characterized as normal-appearing grey matter (NAGM), active and inactive lesions (see Supplementary Figs. 1 and 2).

**Table 2:** List of tissue samples (marked by *) used in this study, showing their control or MS case of origin and the tissue category they contained (CO=control tissue/case; NAGM=normal appearing gray matter). Each line corresponds to a different neocortex or cerebellar cortex tissue block.

| Tissue category | Neocortex |  |  |  | Cerebellar cortex |  |  |  |
| --- | --- | --- | --- | --- | --- | --- | --- | --- |
|  | CO | NAGM | Active Lesion | Inactive Lesion | CO | NAGM | Active Lesion | Inactive Lesion |
| CO1 | * |  |  |  | * |  |  |  |
| CO2 | * |  |  |  |  |  |  |  |
| CO3 | * |  |  |  |  |  |  |  |
|  | * |  |  |  |  |  |  |  |
| CO4 | * |  |  |  | * |  |  |  |
| CO5 | * |  |  |  |  |  |  |  |
|  | * |  |  |  |  |  |  |  |
| CO6 |  |  |  |  | * |  |  |  |
| CO7 |  |  |  |  | * |  |  |  |
| MS1 |  | * |  | * |  |  |  | * |
| MS2 |  | * |  | * |  | * | * |  |
| MS3 |  | * | * | * |  |  | * | * |
|  |  | * |  |  |  |  |  | * |
| MS4 |  |  |  |  |  | * |  |  |
| MS5 |  | * | * | * |  | * | * |  |
| MS6 |  | * |  | * |  |  |  | * |
| MS7 |  | * |  | * |  |  |  | * |
| MS8 |  | * |  | * |  | * |  |  |
|  |  | * |  |  |  |  |  |  |
| MS9 |  |  |  |  |  | * |  |  |
|  |  |  |  |  |  | * |  |  |
| MS10 |  | * | * |  |  |  |  |  |
|  |  | * | * |  |  |  |  |  |
|  |  |  |  | * |  |  |  |  |
| MS11 |  | * |  | * |  |  |  |  |
|  |  | * |  | * |  |  |  |  |
| MS12 |  | * |  | * |  |  |  |  |

### Immunolabeling

#### i) For lesion characterization

Tissue blocks were cut into 20μm thick cryostat sections. Once dried, sections were washed (in Phosphate Buffer Saline - PBS) and fixed in 4% paraformaldehyde (PFA) for 40min. After washing, sections were first incubated for 30min in pure methanol and 2% H_2_O_2_ (for MHC II immunolabeling), or (after immersion for 20min at −20°C in pure ethanol and washes) for 5min in 40% MeOH and 1% H2O2 (for proteolipid protein, PLP, immunolabeling). A blocking step was then performed: 1h with 4% BSA and 0,1% Triton X-100 (for MHC II), or 1h with 10% normal goat serum (Gibco BRL) and 0,4% Triton X-100 (for PLP). Slides were then incubated overnight at 4°C with primary antibodies diluted in the respective blocking solution (for PLP, 0,4% Triton was replaced by 0,2% Triton). After washing, slices were incubated for 45min with biotinylated secondary antibodies (Vectastian kit) diluted in PBS, washed, treated for 1h with the Biotin-Avidin solution (Vectastian kit) and revealed with the DAB chromogen (25mg/50ml), mixed with 25% Tris-base Buffer 1M and H2O2 (4.0E-3% for MHC II or 8.0E-3% for PLP) and Nickel Ammonium Sulfate at 100mg/50ml (for MHC II only). Slides were finally dehydrated into increasing ethanol concentrations (30% to 100%) washed in Xylene and mounted with Eukit. After MHC II immunolabeling, Luxol Fast Blue (LFB) staining was performed before dehydration: sections were immersed in 70% ethanol, incubated overnight with LFB at 60°C, rinsed in 95% ethanol followed by water, and a good contrast was achieved by repeated cycles of incubation in lithium carbonate (0,5g/l), followed by rinses in 70% ethanol and water. Bright field mosaic images of tissue sections labeled for PLP and MHC Class II and LFB were taken with a 5x objective, in order to visualize the whole tissue section and used as references to decide where to focus for AIS analysis acquisitions.

#### ii) For AIS analysis

AIS analysis was performed on sections adjacent to those used for tissue characterization. Dried sections were washed in PBS and fixed for 5min in 2% PFA (in PBS, as for the following incubations). After washing followed by a blocking step in 10% NGS and 0.4% Triton, sections were incubated overnight at 4°C with primary antibodies in 10% NGS and 0.2% Triton. Slides were then washed and incubated for 2h with secondary antibodies diluted in the same solution as for primary antibodies. Finally, slides were washed and mounted with Fluoromount-G (SouthernBiotech). When PLP immunolabeling was added, prior to mounting, a second fixation step (5min in 4% PFA, followed by washing and incubation for 15min at −20°C in pure ethanol) was performed before incubation with the primary and secondary antibodies as before.

### Antibodies

The primary antibodies used were the following: anti-AnkG (mouse IgG2a, Neuromab, clone N106/36, 1:200), anti-PLP (hybridoma of polyclonal rat IgGs, clone AA3, gift from K. Ikenaka, University of Okazaki, Japan, 1:10), anti-Human HLA-DP, DQ, DR (‘anti-MHC II’, mouse IgG1, Dako clone CR3/43, 1:150), anti-Calbindin (mouse IgG1, Sigma-Aldrich, CB-955, 1:500), anti-non-phosphorylated neurofilament H (‘anti-SMI-32’, mouse IgG1, BioLegend clone SMI32, 1:1000). The secondary antibodies used were the following: Alexa 488/Alexa 594 (both 1:1000) /Alexa 647 (1:500) -conjugated secondary antibodies (Invitrogen; Villebon sur Yvette, France) directed against rabbit /rat polyclonal or mouse monoclonal primary antibodies.

### AIS length & soma-AIS gap measurements

AIS analysis was performed by immunolabeling with AnkG and SMI32 (pyramidal neuron marker) in the neocortex or with AnkG and Calbindin (Purkinje cell marker) in the cerebellum. PLP immunolabeling was also performed to further confirm the demyelination status of lesions analyzed. Neocortical SMI32-positive pyramidal neurons belonging to layer 5/6 were identified using hematoxylin-eosin staining on an adjacent section (data not shown), which allowed the identification of the typical large somata of layer 5 pyramidal neurons and the determination of layer 5 extent. Cerebellar Purkinje cells were easily identified with Calbindin immunolabeling. AnkG-immunolabeled AISs from these two neuronal populations were analyzed with an Axiovert 200M (Carl Zeiss) fluorescence microscope, equipped with an Apotome module. Images were acquired with the Axiocam MRm camera (Carl Zeiss) using the Axiovision (Carl Zeiss) image analysis software. Images were taken using 20× (0.8 NA), 40× (1.0 NA), 40× oil (0.75 NA), or 63× oil (1.4 NA) objectives.

AIS length analysis was performed by tracing the AIS in 3D with the Simple Neurite Tracer plugin of Image J (NIH ImageJ software; Bethesda, MD, USA), whereas soma-AIS gap analysis was performed by tracing the soma-AIS gap in 2D with the Image J line tracing tool. AIS length measurements in 3D were restricted to fully intact AISs having clear SMI32 or Calbindin labeling right before and after AnkG labeling or AISs where the typical axon hillock could be clearly observed. The integrity of AISs was further verified in 3D by checking and comparing both channels (AnkG and either SMI32 or Calbindin) in each image plane throughout the volume of the analyzed image and incomplete AISs were not analyzed. As for the soma-AIS gap analysis, we analyzed AISs either affixed to a pyramidal neuron or a Purkinje cell soma or AISs present on a SMI32+ or Calb+ axon connected to its respective soma. Length and soma-AIS gap measurements were performed blinded with respect to the tissue category of origin.

### Statistical analysis

For each AIS, length and soma-AIS gap measures were plotted as individual dots, where each color corresponds to a different tissue block. Both mean and standard error of the mean for each tissue category were calculated. For all these measures (AIS length, soma-AIS gap), to take the intra-individual variability into account, linear mixed-effect models were used, in which the individual factor was considered a random effect. After building the model, a *post-hoc* procedure was applied to assess the difference between each pair of tissue categories, a p-value was therefore computed for each of these pairs and corrected with a Tukey procedure (and reported as such in the results section).

Note that given the atypical distribution of soma-AIS gap values, we recomputed the p-values with permutations (10 000 permutations), followed by a Bonferroni correction for multiple comparisons, and found that it did not change the power of p-values reported (data not shown).

### Model description and simulations protocols

To simulate the neocortical pyramidal cell activity, we used a previously published model available on modelDB (http://modeldb.yale.edu/114394), implemented for the NEURON simulation environment. To simulate the Purkinje cell activity, we used a multi-compartmental model previously developed ^29^, also available on modelDB (http://modeldb.yale.edu/229585) and implemented for the NEURON simulator’s Python interface. Both models are described in supplementary material.

### Data availability

The authors confirm that the data supporting the findings of this study are available within the article [and/or] its supplementary material.

## Results

### Characterization of MS Grey Matter Lesions

To study AISs in MS tissue, we first characterized grey matter lesions found in cortical and cerebellar tissue blocks from MS cases’ brains. Sections were processed with: i) LFB staining (histological myelin stain), to ensure the discrimination between white and grey matter, combined on the same tissue section with MHC II immunolabeling, to detect inflammatory cells; and ii) PLP immunolabeling (myelin marker) on an adjacent section, to allow, better than LFB staining, the discrimination within the grey matter between demyelinated lesions and normally myelinated areas (NAGM and control tissue) (Supplementary Figs. 1 and 2). Tissues were then categorized as either: a) “control” tissue (from non-MS cases), where myelin could be detected both with LFB staining and PLP labeling and no or only few MHC+ cells could be found (Supplementary Figs. 1A and 2A), or (from MS cases) b) “NAGM”, with similar myelinated aspect as in control tissue (Supplementary Figs. 1B, and 2B), c) “active lesion”, where a clear demyelination and a high density of MHC+ cells were detected (Supplementary Figs. 1C and 2C, lesion borders are indicated by dashed lines), d) “inactive lesion”, where clear demyelination was detected, with the absence or a low density of inflammatory MHC+ cells (Supplementary Figs.1D and 2D, lesion border is indicated with a dashed line for the cortical inactive lesion but not for the cerebellar inactive lesion, as the adjacent subregion is devoid of myelin).

### Neocortical layer 5/6 pyramidal neurons have an altered AIS position in both active and inactive MS lesions

We first analyzed the structural properties of AISs known to control the neuron’s spiking properties, namely the length and position of AISs ^30–32^. Given the variability of AIS length between different cortical neuronal subtypes, we focused our analysis on pyramidal neurons (labeled with SMI-32 antibody) belonging to cortical layers 5 and 6 (determined by using hematoxylin-eosin staining). Furthermore, for statistical analyses linear mixed-effect models in which the individual factor was considered a random effect were used, to further eliminate a potential bias due to intra-individual variability (see Materials and Methods).

To ensure the accuracy of (AnkG-labeled) AIS length and position (soma-AIS gap) measurements, the analysis was restricted to fully intact AISs (with SMI-32-positive axon hillock and axonal segments encompassing AnkG labeling, see Materials and Methods for details) but also to AISs connected to their SMI-32 or Calbindin positive cell body. The number of such AISs that could be analyzed was further reduced due to the use of snap-frozen tissue, which not only limited the thickness of sections but also often compromised the integrity of the tissue (hence of AISs).

AIS length and soma-AIS gap were compared between control tissue, MS NAGM, active MS lesions and inactive MS lesions (Fig. 1A-D). As shown on Fig. 1E, the mean length of AISs was: 36.92 ± 0.74μm for control tissue (mean ± standard error of the mean (SEM), n=45 AISs from 5 cases); 39.33 ± 0.62μm for NAGM (n=114 AISs from 14 cases); 40.31 ± 1.60μm for active lesions (n=18 from 2 cases) and 44.27μm ± 1.69μm for inactive lesion (n=33 from 9 cases). No statistically significant difference between any pair of tissue categories was found, except between inactive lesion and NAGM (Fig. 1E).

**Fig. 1:**
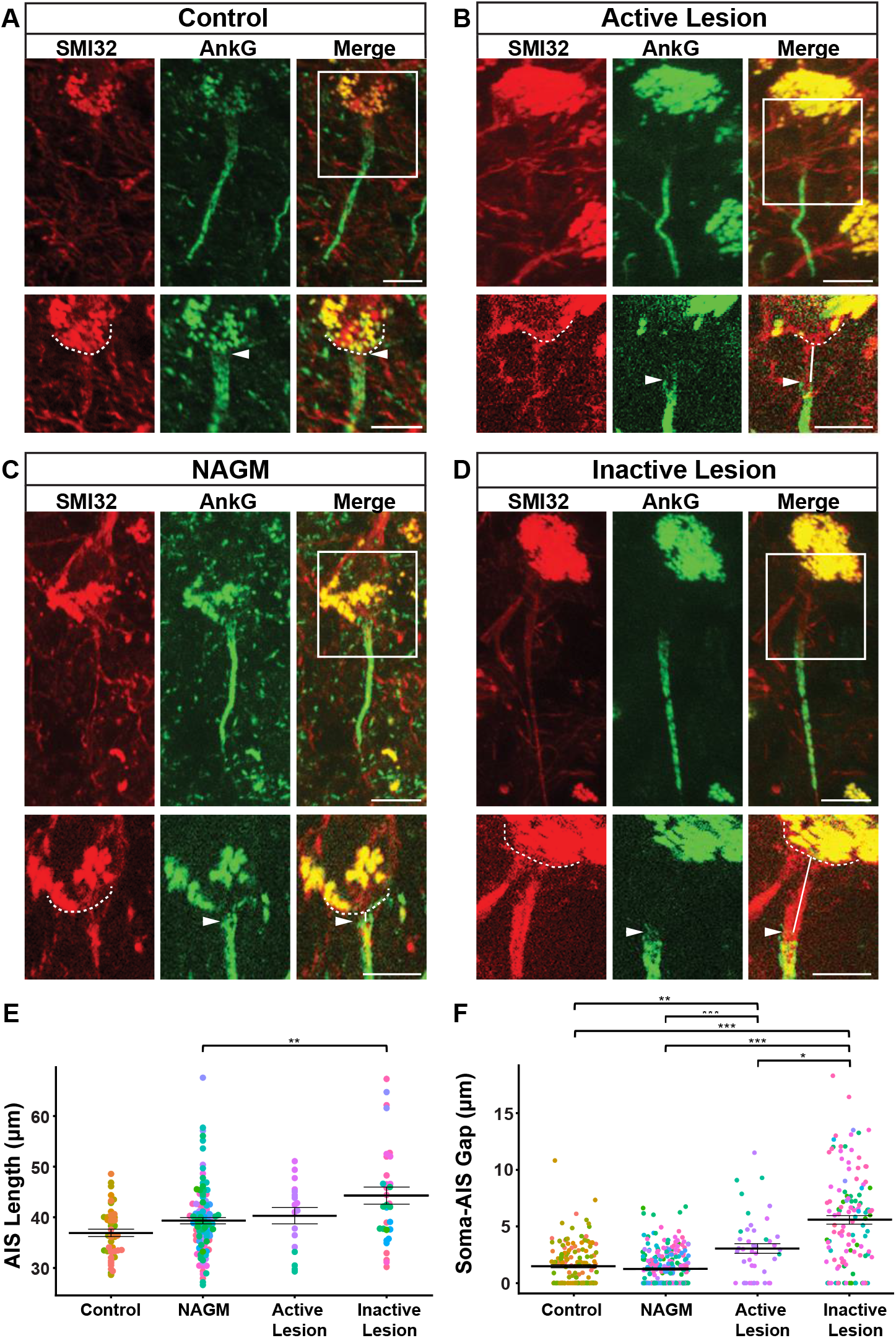
AIS length and soma-AIS gap analysis in neocortical layer 5/6 pyramidal neurons. (A) Representative cortical layer 5/6 pyramidal neurons from control, (B) active MS lesion, (C) NAGM, (D) inactive MS lesion tissues immunolabeled with anti-SMI32 (red), and their AIS immunolabeled with anti-AnkG (green). A higher magnification of the frames indicated with the white rectangles in the merged channel panels are presented in the lower panels for each tissue category. Dashed lines indicate the contour of the soma, arrowheads indicate the start of the AIS, and white lines represent soma-AIS gaps. (E) AIS length (F) Soma-AIS gap measures were plotted in the presented graphs where each dot represents one AIS, and each color represents a different tissue block. For each tissue category, the mean AIS length +/− SEM and the mean soma-AIS gap +/− SEM is represented. A pairwise t-test with a Bonferroni correction for multiple testing was used together with linear mixed effect models to take the intra-individual variability into account. p-values computed for each pair of tissue categories were corrected with a Tukey procedure. Brackets highlight statistically significant differences with * for 0.01 < p < 0.05; ** for 0.001 < p < 0.01; *** for p < 0.001. Scale bar: 10μm.

To analyze whether AIS position was affected in MS tissue, we used the same image acquisitions of AnkG+ AISs from SMI-32+ layer 5/6 pyramidal neurons. SMI-32 labeling allowed to precisely determine the outline of pyramidal neurons somata (dashed lines in Fig. A-D), therefore, to measure the distance between the soma and the beginning of the AIS (pointed by the arrowhead in Fig. 1A-D), the soma-AIS gap. AISs were either directly apposed or very close to the soma in both control tissue and NAGM (mean soma-AIS gap, respectively: 1.48 ± 0.37μm; n= 167 AISs from 5 cases; and 1.24 ± 0.09μm; n= 261 AISs from 9 cases; Fig. 1F). In contrast, the mean soma-AIS gap was significantly increased in both active lesions (3.05 ± 0.42μm; n= 44 AISs from 2 cases) and inactive lesions (5.58 ± 0.37μm; n= 122 AISs from 9 cases), compared to control tissue or NAGM (Fig. 1F). The mean soma-AIS gap in inactive lesions was also significantly increased compared to that in active lesions.

Altogether these results demonstrate that AIS position was altered whereas AIS length was unchanged in both active and inactive demyelinated neocortical lesions.

### Cerebellar Purkinje cells have an altered AIS position in both active and inactive MS lesions

To investigate whether other neuronal populations display similar types of changes in terms of AIS structural characteristics, we analyzed cerebellar Purkinje cells, given their important role in motor coordination and the frequent occurrence of cerebellar grey matter lesions in MS ^3^. Cerebellar control tissue, NAGM, active and inactive lesions were characterized as previously described. Purkinje cells were labeled with an anti-Calbindin antibody and AISs were labeled as described above, allowing Purkinje cell AIS lengths and soma-AIS gaps to be compared between control tissue, MS NAGM, active MS lesions and inactive MS lesions (Fig. 2A-D), using the same approach as the one used for pyramidal neurons.

**Fig. 2:**
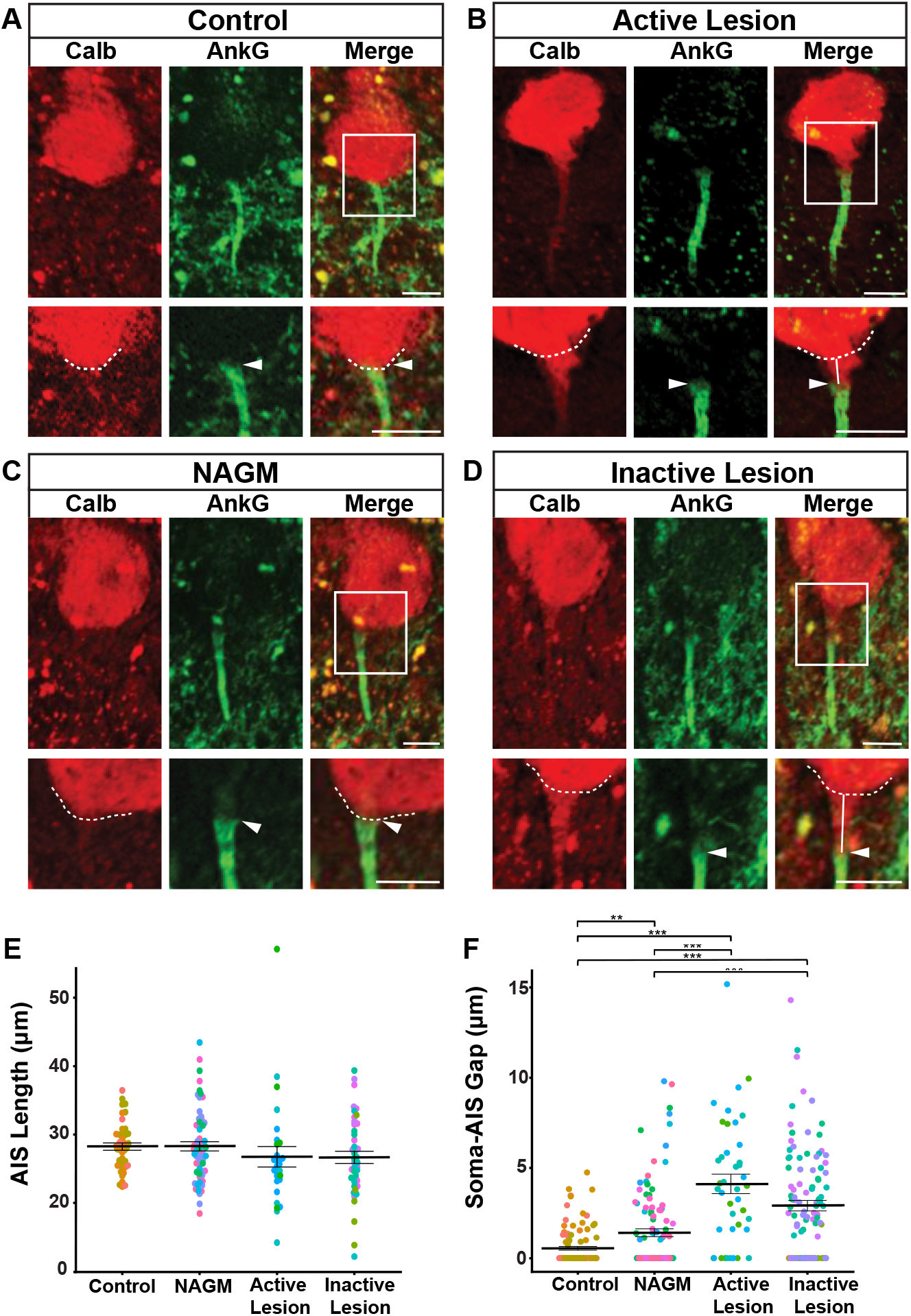
AIS length and soma-AIS gap analysis in cerebellar Purkinje cells. (A) Representative Purkinje cells from control, (B) active MS lesion (C) NAGM (D) inactive MS lesion tissues immunolabeled with Calb and AnkG were presented. A higher magnification of the frames indicated with the white rectangles in the merged channel panels are presented in the lower panels for each tissue category. The dashed line indicates the contour of the cell body, arrowheads indicate the start of the AIS, and white lines represent the soma-AIS gaps. (E) AIS length (F) Soma-AIS gap measures were plotted in the presented graphs where each dot represents one AIS and each color represents a different tissue block. For each tissue category, the mean AIS length +/− SEM and the mean soma-AIS gap +/− SEM is represented. A pairwise t-test with a Bonferroni correction for multiple testing was used together with linear mixed effect models to take the intra-individual variability into account. p-values computed for each pair of tissue categories were corrected with a Tukey procedure. Brackets highlight statistically significant differences with * for 0.01 < p < 0.05; ** for 0.001 < p < 0.01; *** for p < 0.001. Scale bar: 10μm.

As shown on Fig. 2E, the mean length of AISs was: 28.23 ± 0.54μm for control tissue (n= 43 AISs from 4 cases); 28.28 ± 0.67μm for NAGM (n=62 AISs from 5 cases); 26.74 ± 1.50μm for active lesions (n=28 AISs from 3 cases); and 26.66 ± 0.90μm for inactive lesion (n=43 AISs from 4 cases). No statistically significant difference between any pair of tissue categories was found (Fig. 2E).

To analyze whether AIS position was affected in MS tissue, the same image acquisitions of AnkG+ AISs from Calb+ Purkinje cells were used and soma-AIS gap was measured. Calbindin labeling allowed to precisely determine the outline of Purkinje cells somata (dashed lines on Fig. 2A-D), therefore, to measure the distance between the soma and the beginning of the AIS (pointed by the arrowhead in Fig. 2A-D), the soma-AIS gap. Whereas AISs were almost affixed to the soma in control tissue (mean soma-AIS gap: 0.55±0.10μm, n= 100 AISs from 4 cases), a short gap was observed in NAGM (mean soma-AIS gap: 1.41 ± 0.22μm, n= 110 AISs from 5 cases; Fig. 2F). In contrast, a significant increase of soma-AIS gap was evidenced in both active lesions (4.12 ± 0.54μm, n= 41 AISs from 3 cases) and inactive lesions (2.91 ± 0.29μm; n= 105 AISs from 4 cases) compared to control tissue or NAGM (Fig. 2F).

Altogether these results show that AIS position but not length was altered in both active and inactive cerebellar demyelinated lesions.

Taken together, this morphological analysis of AISs from neocortical layer 5/6 pyramidal neurons and from cerebellar Purkinje cells shows that compared to control tissue AIS length is unchanged within active and inactive demyelinated lesions and NAGM. In contrast, AIS position is altered within active and inactive demyelinated MS lesions, with an AIS being shifted away from the soma, a change that may lead to altered excitability properties.

### Functional consequences of AIS changes occurring in active demyelinated MS lesions, assessed by computational modeling

Biophysical analyses of spike initiation in neurons highlighted the role of AIS geometry on neuronal excitability ^30,33^. To assess the potential effect of the increased soma-AIS gap observed in active and inactive demyelinated MS lesions, we simulated and analyzed the excitability properties of both a model neocortical pyramidal cell (Fig. 3A) and a model Purkinje cell (Fig. 3B), based on detailed morphological reconstructions ^29,34^.

**Fig. 3:**
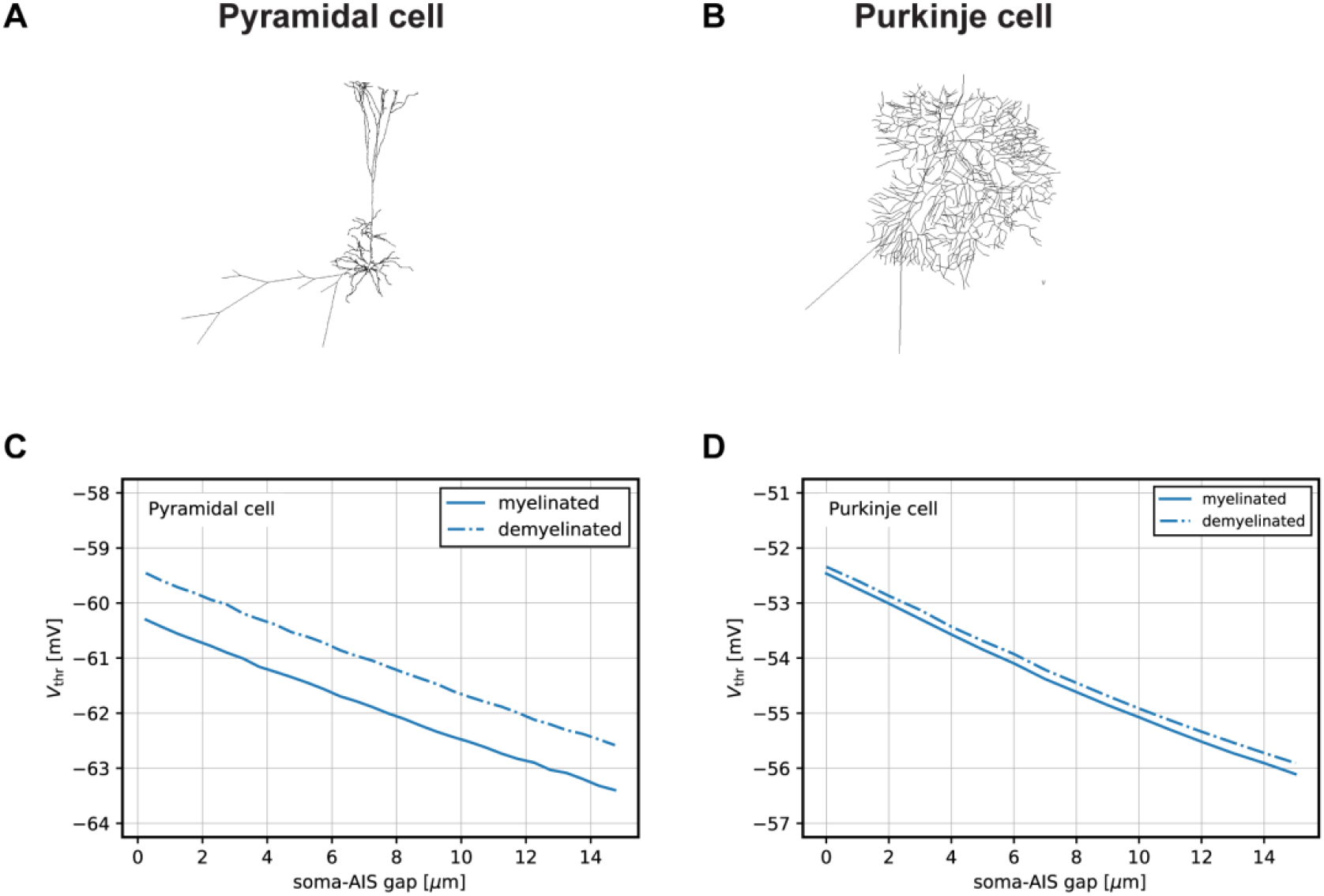
Modelling of the functional consequences of AIS-soma gap changes observed in active demyelinated MS lesions in pyramidal and prkinje cells. (A) Morphology sketch of pyramidal model cell. (B) morphology sketch of Purkinje model cell. (C), Variation of the AP voltage threshold with the length of the soma-AIS gap in pyramidal model cell. (D) Variation of the AP voltage threshold with the length of the soma-AIS gap in Purkinje model cell.

For the pyramidal cell, we found that the AP voltage threshold decreases with increasing soma-AIS gap (−0.28mV/µm, Fig. 3C, solid line). This effect persists and is comparable when the demyelinated status of the axon is taken into account (Fig. 3C, dashed line), which depolarizes the threshold by about 0.8mV.

For the Purkinje cell, we similarly found that the AP voltage threshold decreases with increasing soma-AIS gap (−0.25 mV/µm, Fig. 3D, solid line), while demyelination alone leads to a very minor (~0.1mV) depolarization of the threshold (Fig. 3D, dashed line).

Overall, these results show that the increased soma-AIS gap can cause AP voltage threshold decrease in both neocortical pyramidal neurons and Purkinje cells, consistent with theoretical predictions for neurons in the ‘resistive coupling regime’ ^30,33^. Under the assumption that other parameters that might affect spike threshold do not change, the observed changes in the soma-AIS gap may thus cause both of these neurons to be more excitable, i.e., more prone to spike, than in control tissue.

## Discussion

Grey matter damage is now established as an important contributor to long-term disability in MS. Grey matter atrophy results from a combination of demyelination, neurite transection and loss, as well as neuronal loss, partially related to inflammation ^35^. In addition, other alterations at the synaptic level have been described, such as reduction in synaptic density ^7,36^ or dendritic spine loss ^8^. Although reduced synaptic density was mainly reported within cortical demyelinated lesions and the role of both demyelination and inflammation has been suggested, dendritic spine loss has been found both in lesions and NAGM, favoring the idea of a diffuse alteration partially independent of demyelination.

Despite its major role in neuronal function, whether the AIS is affected in MS had not been addressed. In this context, AIS alterations have been reported in both cuprizone ^27,28^ and EAE mouse models of MS ^27^. Here we demonstrate for the first time that in MS the distance between soma and AIS is increased, for both neocortical pyramidal neurons and cerebellar Purkinje cells, in active as well as inactive demyelinated lesions, while the length of their AIS is unaffected. In addition, we provide computational evidence that this increased soma-AIS gap could lead to a decreased AP voltage threshold, which would make these neurons more prone to spike.

This study of AISs in MS post-mortem tissue was challenging as several factors reduced the population size of AISs analyzed. We overcame the difficulty of immunolabeling AISs on paraffin-embedded sections by working on snap-frozen samples, where immunolabeling was reliable and reproducible. However, tissue quality and limitation of section thickness (the best results were obtained with 20 µm) reduced the number of fully intact AISs particularly for the 3D length analysis. In addition, obtaining cortical samples containing active MS lesions (from early stages of the disease) was another challenge but we analyzed more sections from such lesions to increase the number of fully intact AISs. As we included some distally cut (i.e., by cryostat sectioning) AISs to our 2D gap analysis (as long as the axon hillock and its corresponding cell body were clearly identifiable with either the SMI32 or the Calbindin labeling), more AIS-soma gap than AIS length measurements could be performed in each tissue category.

Finally, despite the limited population size of AISs that could be analyzed, the use of a powerful statistical analysis (with linear mixed effect models) to take the intra-individual variability into account, allowed us to demonstrate significant differences with respect to structural characteristics of AISs, in MS lesions compared to control tissue.

Very similar AIS phenotypes (unaltered AIS length but altered AIS location) were found in both neocortical pyramidal neurons and Purkinje cells in MS grey matter demyelinated lesions, suggesting that a common molecular mechanism may be at play in both cell types. The AIS has been found to be altered in different ways in many pathological conditions ^37–39^. Yet, the mechanisms responsible for these changes remain largely elusive. On the one hand, a calpain-dependent mechanism has been shown to cause AIS dismantling through AIS constituent protein degradation ^25^. On the other hand, a calcium- and calmodulin-dependent protein phosphatase calcineurin has been shown to mediate changes in soma-AIS gap, with the involvement of phosphorylated myosin light chain, an activator of contractile myosin II^40,41^. However, the mechanisms responsible for calcineurin activation and for mediating calcineurin activity still need to be elucidated.

The fact that we observed these alterations in both active and inactive demyelinated MS lesions suggests that it is not a late feature of chronically demyelinated axons but that it might occur early in the lesion process. The detection of longer AIS-soma gaps in active and even more profoundly in inactive grey matter lesions also suggests that, even though both inflammation and demyelination are involved in the increased AIS-soma gap phenotype, demyelination per se is likely to play a more prominent role. Therefore, further studies are required to shed a light into the mechanisms by which AISs are altered in MS.

Changes in the soma-AIS gap can have a number of opposing effects on spike initiation, depending on the characteristics of the soma and the AIS ^30^. Biophysical analyses and modeling suggested that small displacements of an AIS with a fixed number of Nav channels generally cause a decreased AP threshold as the soma-AIS gap increases ^33,42,43^ in accordance with results obtained in neocortical pyramidal cells showing that a decrease in soma-AIS gap depolarized AP threshold ^28^. In order to quantitatively assess the effects of the soma-AIS gap changes in MS lesions, and to take the demyelination of MS lesions into account, we simulated computational models of a pyramidal cell and a Purkinje cell based detailed morphological reconstructions.

The overall effect of the modeled AISs changes in both pyramidal neurons and Purkinje cells showed a slight decrease in AP voltage threshold, and we found that demyelination per se did not have a profound effect. These small changes could nevertheless alter the relationship between synaptic input and cell firing and potentially lead to neurological symptoms. However, we assumed here that other parameters such as Nav and Kv channel densities in the AIS remain unchanged. Further studies of potential modifications of Navs and Kvs in MS active and inactive lesions are certainly required to better understand how neuronal function is affected in addition to the effects caused by the structural change in AIS we detected in the current study (Fig. 4).

**Fig.4:**
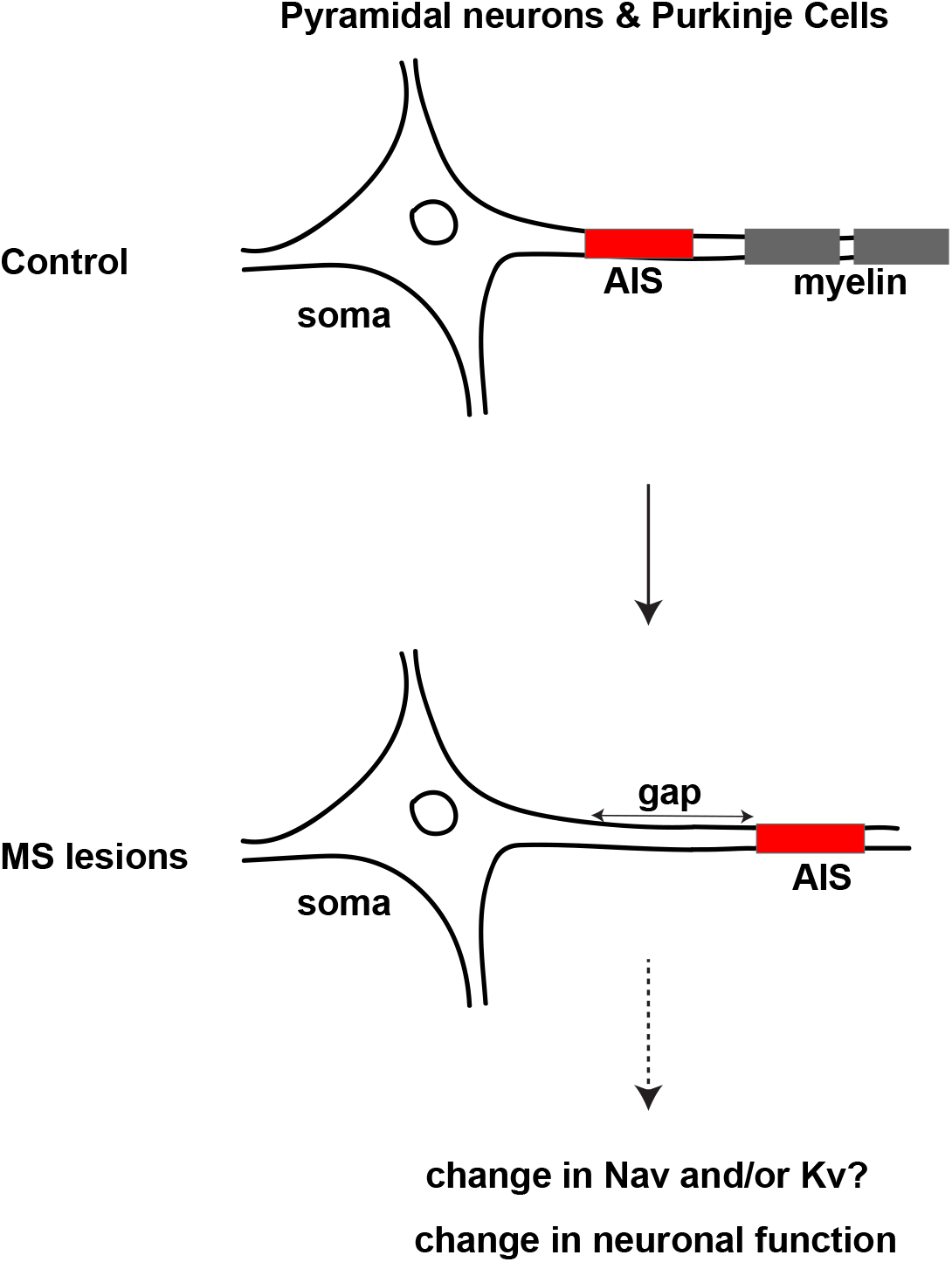
Schematic presentation of effect of AIS changes in MS tissue. Whilst the AIS length remained unchanged, the AIS-soma gap was found to be increased in demyelinated active and inactive MS lesions compared to control tissue, which, together with potential accompanying changes in Navs and Kvs, may alter neuronal function.

## Conclusion

The frequency and functional impact of MS grey matter lesions on disease progression is now well established ^6,35^. Here we show for the first time that the AIS position is affected in MS lesions, and that this change could play a role in functional alterations. These findings highlight the fact that not only nodes of Ranvier but also AISs can contribute to the neurodegeneration observed in MS. Finally, the mechanisms of how and when AIS alteration may contribute to the disease development and progression remain to be elucidated.

## Abbreviations

AIS: axon initial segment
AnkG: ankyrinG
AP: action potential
EAE: experimental allergic encephalomyelitis
Kv: voltage-gated potassium channels
LFB: luxol fast blue
MHC: major histocompatibility complex
MOG: myelin oligodendrocyte glycoprotein
MS: Multiple Sclerosis
NAGM: normal-appearing grey matter
Nav: voltage-gated sodium channels
PLP: proteolipid protein

## Acknowledgments

Part of this work was carried out on the Histomics and icm.Quant core facilities. We gratefully acknowledge Annick Prigent and Brigitte Zeau for their technical help in handling human tissues, and Aurélien Dauphin, Dominique Langui and Claire Lovo for their help in image acquisitions and quantification of AISs is in 2D and 3D. We thank Ivan Moszer for statistical analysis discussions.

## SUPPLEMENTARY MATERIAL

### Model description

#### Neocortical pyramidal cell

We made the following minor modifications to the neocortical pyramidal cell model ^34^ for the sake of simplicity: we approximated the AIS geometry to three linear segments (that closely follow the AIS approximately piecewise linear decrease in diameter), defined by the four following diameters along the AIS: d(l0) = 5 µm, d(l_1_) = 2 µm, d(l_2_) = 1.7 µm, and d(l_3_) = 1.6 µm, with l_0_ = 0 µm (proximal end of the AIS), l_1_ = 0.55 l_AIS_, l_2_ = 0.95 l_AIS_, and l_3_ = l_AIS_ (distal end of the AIS).

The model includes a single Nav channel and several types of Kv channels with Hodgkin-Huxley-like kinetics. The main parameters are as follows (please refer to ^34^ for additional detail): E_Na_ = 55 mV, g_Nav;D/S_ = 60 pS/µm^2^ in dendrites and soma and g_Nav;NoR_ = 2500 pS/µm^2^ in nodes of Ranvier (intermodal distance: 60 µm). Three types of Kv channels describe (i) a high-voltage-activated channel (“Kv”), (ii) a faster low-voltage activated Kv1-like channel (“Kv1”), and (iii) a slowly-activating and non-inactivating M-type channel (“Km”). E_K_ = –85 mV. Peak conductances (identical for dendrites and soma) are: g_Kv;D/S_ = 20 pS/µm^2^, g_Kv1;D/S_ = 100 pS/µm^2^, and g_Km;D/S_ = 5 pS/µm^2^. In order to avoid hypotheses about the precise spatial distribution of ion channels in the AIS, we distributed the identical number of ion channels as in the original model on the simplified AIS (cone + tube), with constant density along the AIS: g_Nav;AIS_ = 2917 pS/µm^2^, g_Kv;AIS_ = 86 pS/µm^2^, g_Kv1;AIS_ = 161 pS/µm^2^, and g_Km;AIS_ = 6.3 pS/µm^2^. g_Kx;NoR_ = g_Kx;AIS_. A hyperpolarization-activated current is implemented by I_h_ channels distributed in the soma and dendrites with an exponential increase in density with distance from the soma. The passive electrical properties of the reconstructed cell were set as follows: C_m_ = 0.9 µF/cm^2^, R_i_ = 100 Ωcm, and R_m_ = 15 kΩcm^2^.

##### Purkinje cell

In the Purkinje cell model ^29^, the geometry of the AIS is not based on a morphological reconstruction but represented by a single segment of constant diameter. We did the following minor modifications to the AIS geometry: while keeping the original AIS length (21 µm), we changed the AIS diameter (to 1.94 µm) to accommodate for a larger fraction of Nav channels at the AIS.

The model contains Na^+^, K^+^, and Ca^2+^ conductances with several subtypes for Kv, Ca-dependent potassium (Kca), and calcium (Ca) channels based on Markovian or Hodgkin-Huxley-like dynamics ^29^. It furthermore contains a mixed cationic channel (HCN1) as well as explicit dynamics for the internal calcium buffer ^29^. The model for the Nav channel is based on a Markovian state dynamics and accounts for transient, persistent and resurgent Na^+^-current components ^44^. E_Na_ = 75 mV. We kept the same total number of Nav channels as in the original model but distributed them slightly differently to reduce the ratio of somatic to AIS Nav channel density. Peak conductances were: g_Nav;D_ = 160 pS/µm^2^ in dendrites, g_Nav;S_ = 1861 pS/µm^2^ in the soma and g_Nav;AIS_ = 8095 pS/µm^2^ in the AIS (g_Nav;S_/g_Nav;AIS_ ~ 0.23 as opposed to ~0.43 in the original model); g_Nav;NoR_ = 300 pS/µm^2^ in nodes of Ranvier (internodal distance: 100 µm). We furthermore distributed the AIS Kv channels homogenously along the AIS while keeping the average channel density at the AIS constant, leading to g_Nav;AIS_ = 3948 pS/µm^2^, g_Kv1.1;AIS_ = 12 pS/µm^2^, and g_Kv3.4;AIS_ = 49 pS/µm^2^. All other parameters were unchanged. Passive electrical properties were set as follows: C_m_ = 0.77 µF/cm^2^, R_i_ = 122 Ωcm, and R_m_ = 0.91 kΩcm^2^. For all other parameters, please refer to ^29^.

#### Simulation protocols

##### Modification of AIS morphology

We systematically varied the soma-AIS gap and AIS channel densities as follows. To mimic a finite soma-AIS gap, of length l_gap_, we set channel densities along this soma-AIS gap to their somatic values and accordingly extended the AIS distally at constant diameter on a length l_gap_ to preserve the total AIS length.

##### Myelination and demyelination (pyramidal cell)

Kole et al. originally mimicked myelination by reducing C_m_ to 0.02 µF/cm^2^ along the internodal sections^34^. Since effective channel conductances might have to reflect the high resistance imposed by a myelin sheath, we decided to additionally suppress ion channels in myelinated internodes (setting peak conductance values to zero). Conversely, demyelination was mimicked by restoring the internodal membrane capacitance to the default value C_m_ = 0.9 µF/cm^2^ and by setting channel peak conductances to finite values that were determined as follows. Based on the observation that ion channels are redistributed in the membrane of axons upon disorganized nodes of Ranvier in demyelinated MS lesions ^10,11^, we considered peak conductances of demyelinated internodes and within the nodes to be given by the approximate mean value of the respective somatic and node peak conductances: g_Nav;dem_ = 106 pS/µm^2^, g_Kv;dem_ = 41 pS/µm^2^, g_Kv1; dem_ = 22 pS/µm^2^, and g_Km; dem_ = 5.5 pS/µm^2^.

##### Myelination and demyelination (Purkinje cell)

In line with the original model, we mimicked myelination by a vanishing C_m_ and vanishing ionic conductances along the internode. In our study, we mimicked demyelination by restoring the internodal membrane capacitance to the default value C_m_ = 0.77 µF/cm^2^. We furthermore reduced channel concentrations within the nodes to the average densities between internodes and nodes in myelinated conditions. For Nav, we thus obtained g_Nav;dem_ = 13 pS/µm^2^.

##### Determination of the voltage threshold

Because we were mainly interested in the impact of AIS morphological parameters on the voltage threshold, the procedure used to determine a voltage threshold *V*_thr_ for APs recorded in the soma is the following: whenever the time derivative of the somatic voltage d*V*(*t*)/d*t* crosses a fixed threshold *c*, the voltage threshold is given by the somatic potential at the time *t_c_* of threshold crossing, *V*_thr_ = *V*(*t_c_*). In the case of multiple threshold crossings, *V*_thr_ did not vary between APs. While the absolute values of *V*_thr_ depend slightly on the choice of *c*, the relative variation does not, nor did the number of threshold crossings within a reasonable range. Throughout this study, we used *c* = 20 mV/ms.

##### Current injection

To elicit stationary firing in the pyramidal cell, a constant current of 0.9 nA was injected in the soma during the simulation. The Purkinje cell was spontaneously active, thus no current was injected.

### Supplementary Figure Legends

**Supplementary Fig. 1:**
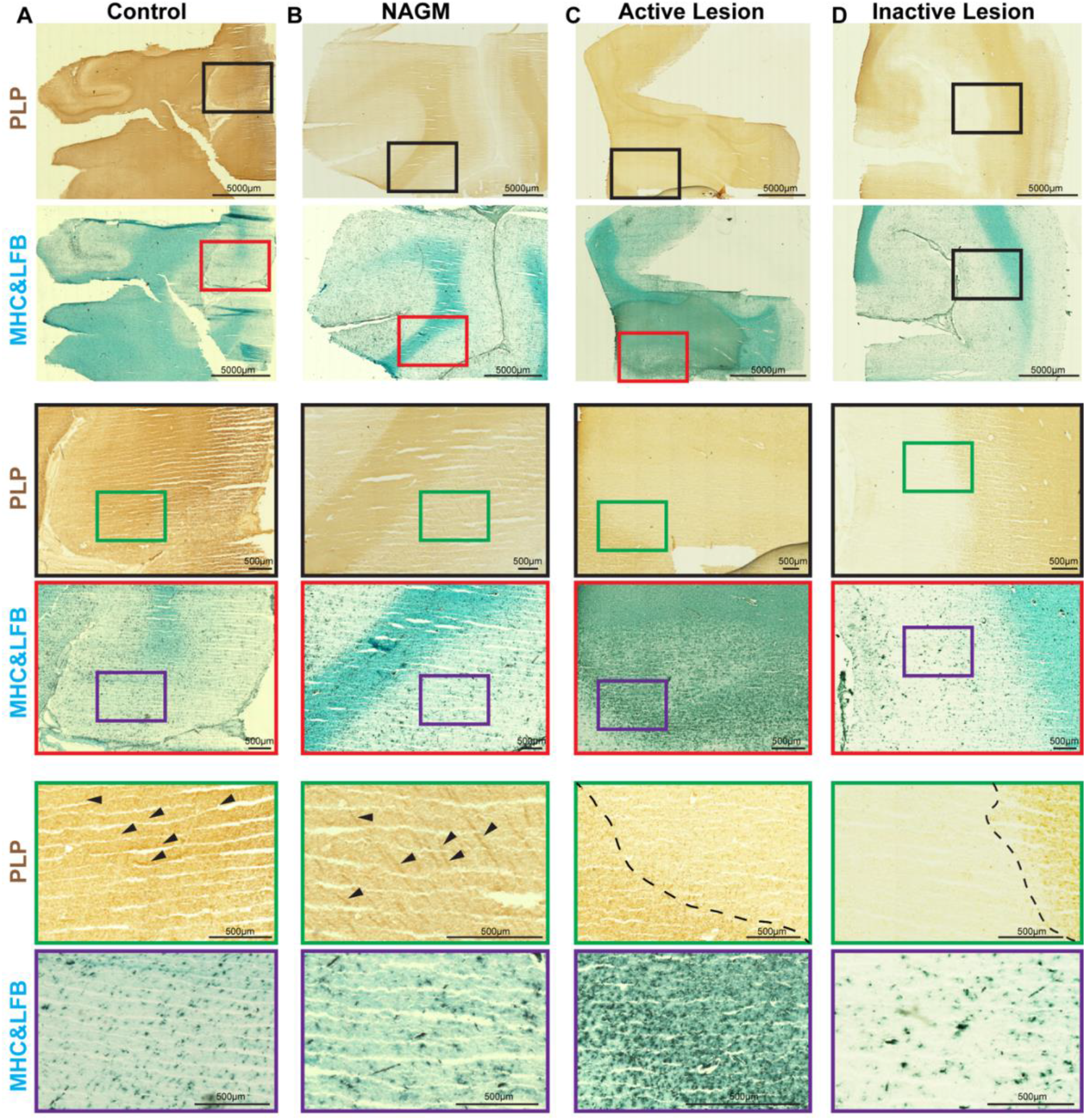
Characterization of Cortical MS lesions. PLP immunolabeling and LFB staining combined with MHC immunolabeling of (A) a control section (B) a NAGM section (C) an active lesion section (D) an inactive lesion. Higher magnification of the selected regions from PLP (indicated by the black frames) and LFB & MHC (indicated by the red frames) are presented. Further magnification of the selected sub-regions from these frames, indicated by the green and purple frames for PLP and LFB & MHC labelings’ respectively are also presented for better visualization. Arrowheads show the “stripe-like” myelin fibers labeled with PLP in control and NAGM panels and dashed lines represent the lesion borders in active and inactive lesion panels. Scale bars: 5000µm for mosaic images and 500µm for zoomed images.

**Supplementary Fig. 2:**
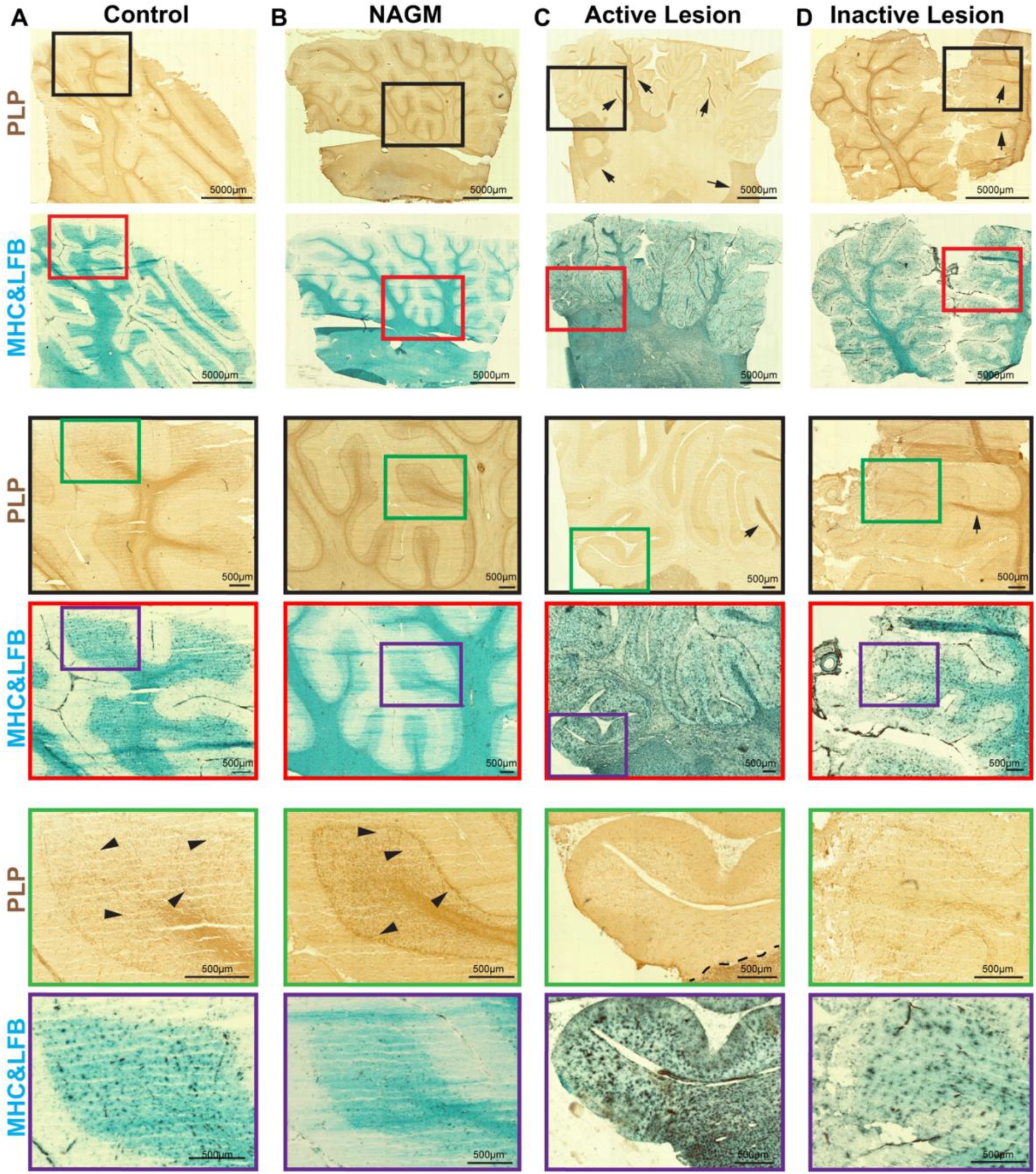
Characterization of Cerebellar MS lesions. PLP and LFB & MHC labeling of (A) a control section (B) a NAGM section (C) an active lesion section (D) an inactive lesion section is presented. Higher magnification of the selected regions from PLP (indicated by the black frames) and LFB & MHC (indicated by the red frames) are presented. Further magnification of the selected sub-regions from these frames, indicated by the green and purple frames for PLP and LFB & MHC labelings’ respectively are also presented for better visualization. Arrowheads show the “stripe-like” myelin fibers labeled with PLP in control and NAGM panels, arrows show the remaining myelinated white matter tracts in almost fully demyelinated selected regions, dashed line represent the lesion border in the active lesion panel while no border is represented for inactive lesion panel as the selected area is fully demyelinated. Scale bars: 5000µm for mosaic images and 500µm for zoomed images.

## Abbreviated Summary

Dilsizoglu Senol et al. highlight the importance of axon initial segment in multiple sclerosis by demonstrating its shift from the soma in both neocortex and cerebellum of multiple sclerosis cases having active and/or inactive grey matter lesions and by showing evidence that this change could affect neuronal function.

